# FetoML: Interpretable predictions of the fetotoxicity of drugs based on machine learning approaches

**DOI:** 10.1101/2023.09.27.559678

**Authors:** Myeonghyeon Jeong, Sunyong Yoo

**Affiliations:** Department of ICT Convergence System Engineering, Chonnam National University, Gwangju, Republic of Korea

## Abstract

Pregnant females may use medications to manage health problems that develop during pregnancy or that they had prior to pregnancy. However, using medications during pregnancy has a potential risk to the fetus. Assessing the fetotoxicity of drugs is essential to ensure safe treatments, but the current process is challenged by ethical issues, time, and cost. Therefore, the need for *in silico* models to efficiently assess the fetotoxicity of drugs has recently emerged. Previous studies have proposed successful machine learning models for fetotoxicity prediction and even suggest molecular substructures that are possibly associated with fetotoxicity risks or protective effects. However, the interpretation of the decisions of the models on fetotoxicity prediction for each drug is still insufficient. This study constructed machine learning-based models that can predict the fetotoxicity of drugs while providing explanations for the decisions. For this, permutation feature importance was used to identify the general features that the model made significant in predicting the fetotoxicity of drugs. In addition, features associated with fetotoxicity for each drug were analyzed using the attention mechanism. The predictive performance of all the constructed models was significantly high (AUROC: 0.854–0.974, AUPR: 0.890–0.975). Furthermore, we conducted literature reviews on the predicted important features and found that they were highly associated with fetotoxicity. We expect that our model will benefit fetotoxicity research by providing an evaluation of fetotoxicity risk for drugs or drug candidates, along with an interpretation of that prediction.

**Author summary:** Drugs are often necessary for the treatment of diseases in pregnant females. However, some drugs can potentially cause fetotoxicities, such as teratogenicity and abortion. Therefore, it is essential to study fetotoxicity, but traditional toxicity testing demands time, money, and labor. To modernize these testing methods, *in silico* approaches for predicting the fetotoxicity of drugs are emerging. The proposed models so far have successfully predicted the fetotoxicity of drugs and proposed some fetotoxicity-related substructures, but the interpretation of the model’s determination is still insufficient. In this study, we proposed FetoML to predict the fetotoxicity of drugs based on machine learning and provide the substructures that the model focused on in predicting fetotoxicity for each drug. We confirmed the significant predictive performance and interpretability of the model through a quantitative performance evaluation and literature review. We expect FetoML to benefit fetotoxicity studies of drugs by modernizing the paradigm of fetotoxicity testing and providing insights to researchers.

## Introduction

Taking medications during pregnancy carries potential fetotoxicity risks, including congenital disabilities, in-utero death, and growth retardation [1]. However, treatment with medication may be essential for the health of the pregnant female and the normal development of the fetus. In a web-based survey of medication use during pregnancy with multinational study settings in Europe (Western, Northern, and Eastern), North and South America, and Australia, 81.2% of pregnant females reported using at least one medication [2]. Surveys of medication use during pregnancy conducted in various countries have also reported similar findings [3–8]. Despite the common use of drugs during pregnancy, the fetotoxicity risks of many drugs are unknown [9]. These results may be attributed to the exclusion of pregnant females from clinical trials, which is often based on ethical concerns regarding potential risks to fetal and infant health. The fetotoxicity risk of drugs in human pregnancy was assessed through the results of exposure cohort studies and large population-based case-control studies. While such post-marketing surveillance is essential to assess fetotoxicity in humans, it is a retrospective approach that addresses the problem after the drug has been exposed to pregnant females and their fetuses, which can lead to serious public health concerns. Therefore, fetotoxicity assessments in human pregnancies are performed through nonclinical testing before the approval of drugs. However, testing paradigms involving developmental and reproductive toxicology (DART) studies in mammalian species have not changed significantly since the 1960s [10], and pharmacodynamics and pharmacokinetic differences between species do not clearly explain human fetotoxicity. Additionally, DART studies on rodents and rabbits require between 16 to 20 litters [11], which is ethically problematic, time-consuming, and has high costs. Therefore, there is an emerging need for *in silico* models to modernize the testing paradigm and reduce animal use, costs, and time to assess the fetotoxicity of drugs [10, 12]. This implies that *in silico* models can offer promising solutions to the challenges of the current fetotoxicity assessment.

*In silico* toxicity assessments leverage the structure and properties of compounds using computational models to predict and analyze various toxicities and side effects [13]. These allow for the prioritization of samples that need to be tested *in vitro* or *in vivo*, which can minimize the cost and the number of animals used in testing. *In silico* toxicology is performed using expert methods, such as structural alerts and read-across, and machine learning-based approaches, such as ligand-based or structure-based approaches [14]. Machine learning-based approaches have been recently promoted as they have shown high performance with large amounts of data. Although machine learning-based drug toxicity predictions have been successful, most of the machine learning models are ‘black box,’ meaning that they lack explanations of how they made inferences. Providing explanations of the model’s inferences improve trust and transparency of the model while also providing valuable insights for practitioners [15]. For this reason, the field of eXplainable AI (XAI) is rapidly developing, with various algorithms such as self-attention (intra-attention) [16], Local Interpretable Model-Agnostic Explanations (LIME) [17], and Shapley Additive Explanations (SHAP) [18]. These algorithms can explain the reasoning behind the model predictions, increasing interpretability and transparency. In particular, it is crucial in toxicity assessment to provide practitioners with a better understanding of compounds’ mechanisms of action [19]. Recent studies have confirmed that XAI can be effectively utilized in machine learning-based toxicity prediction models [20, 21].

Motivated by the need for *in silico* models for fetotoxicity assessment, several approaches have been previously proposed. Some approaches have used pharmaceutical claims databases or Electronic Health Record (EHR) data to identify whether medications affect the fetus [22, 23]. However, these approaches cannot predict the risk of fetotoxicity prior to market approval of the drug since they use data that is unavailable for preclinical trials. The other approaches are machine learning-based models that use datasets containing fetotoxicity information for drugs [24–26]. These models use the structural, physiological, or physicochemical features of the drug as the input to predict the fetotoxicity. These proposed approaches have been shown to significantly predict the fetotoxicity of drugs and have identified specific features associated with fetotoxicity. However, their predictions do not provide an explanation of which specific features of each drug the model focused on to make its decision. Providing researchers with an explanation of which drug features were focused on when making the predictions can provide important insights into the fetotoxicity test design.

In this study, we constructed models for predicting the fetotoxicity of drugs and drug candidates based on various machine-learning approaches. Our models provide predictions of the fetotoxicity risk with explanations for the decisions. The self-attention neural network (NN) approach is interpretable in terms of the specific features that were focused on when making decisions for each drug. It can provide insights into the fetotoxicity of the given compounds at the drug development stage or studies of fetotoxicity. The machine learning models, excluding the self-attention NN, used the permutation feature importance method to identify the features that were important to the prediction of the model.

## Results

### Performance evaluation

In this study, we constructed various machine learning models based on logistic regression (LR), support vector machine (SVM), random forest (RF), extra trees (ET), gradient boosting machine (GBM), extreme gradient boosting (XGBoost), and self-attention NN algorithms to predict fetotoxicity. We performed the quantitative performance evaluation to assess the ability of all the constructed models to predict fetotoxicity. In addition, we compared the performance of each model to determine which machine learning algorithm had the best outcome. First, the receiver operating characteristic (ROC) and precision-recall (PR) curves of each model are presented in Fig 1. All the models had significant prediction performance (area under the ROC (AUROC): 0.854–0.974, area under the PR (AUPR): 0.890–0.975), in contrast, the completely randomized classification model had an AUROC and AUPR close to 0.5. The ET-based model had the highest AUROC (0.974) and AUPR (0.975). Along with this, we evaluated the following performance metrics: accuracy, precision, recall, and F1-score. The threshold for each model was set to the value with the highest F1-score. All the performance metrics evaluated for each model are shown in Table 1. The model with the highest accuracy, recall, and F1-score was the SVM-based model. The LR-based and ET-based models had the best performance for precision and recall, respectively.

**Fig 1.**
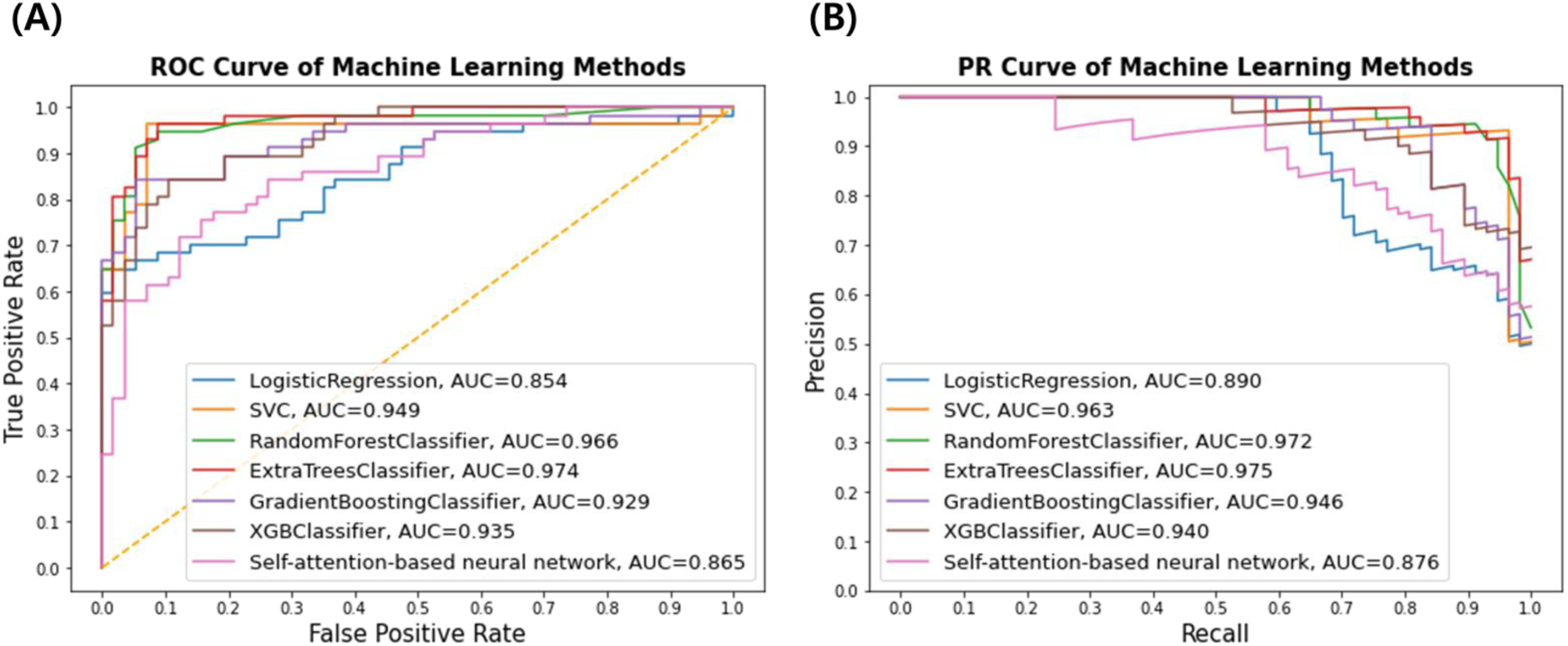
ROC and PR curves for each machine learning model. (A) ROC curve plotting the change in the true positive rate (TPR) and false positive rate (FPR) in response to a change in threshold. The models with better performance are more curved to the upper left. (B) PR curve plotting the change in precision and recall in response to a change in threshold. The models with better performance are more curved to the upper right.

**Table 1.** Evaluation results for all the performance metrics in each model. The LR model achieved the highest precision metric, the SVM model demonstrated the highest accuracy, recall, and F1-score, while the ET model exhibited the highest metrics for AUROC, AUPR, and recall.

| Model | Accuracy | AUROC | AUPR | Precision | Recall | F1-score |
| --- | --- | --- | --- | --- | --- | --- |
| LR | 0.816 | 0.854 | 0.890 | **0.974** | 0.649 | 0.779 |
| SVM | **0.947** | 0.949 | 0.963 | 0.932 | **0.965** | **0.948** |
| RF | 0.930 | 0.966 | 0.972 | 0.915 | 0.947 | 0.931 |
| ET | 0.939 | **0.974** | **0.975** | 0.917 | **0.965** | 0.940 |
| GBM | 0.895 | 0.929 | 0.946 | 0.941 | 0.842 | 0.889 |
| XGBoost | 0.868 | 0.935 | 0.940 | 0.889 | 0.842 | 0.865 |
| Self-attention NN | 0.842 | 0.857 | 0.906 | 0.868 | 0.807 | 0.836 |

### Analysis of model predictions

In the collected dataset, the fetotoxicity of a drug is determined by the observed reports of fetotoxic development from pregnant females taking that drug. Therefore, if a drug is actually fetotoxic in humans but has not been reported, it can be classified as having ‘non-fetotoxicity.’ As such, despite the mislabeling of some of the data in the test set, the model was able to predict the actual value. We analyzed the false positive results of the model to identify drugs that were not reported as ‘fetotoxic.’ The false positive results with high prediction scores for each model are presented in Table 2. The drug ‘brimonidine tartrate,’ predicted as ‘fetotoxic’ by most of the models among the false positive drugs, is used to treat glaucoma. The drug was labeled as ‘non-fetotoxic’ since it was classified as ‘category B3’ in the TGA dataset. Brimonidine tartrate is an eye drop commonly used to lower short- and long-term intraocular pressure in patients with glaucoma [27]. No well-controlled human studies have been conducted on the teratogenicity of the drug [28]. However, the drug may cross the placenta and reach the human fetus, across the blood-brain barrier, and cause central nervous depression and apnea [29]. This means that there are no reports of toxicity to the human fetus, but there is a high potential risk of fetotoxicity. Also, ‘decitabine’ is a drug with a high prediction score for fetotoxicity in the RF, ET, and GBM models. It was labeled as ‘non-fetotoxic’ according to the ‘2nd class’ in the KIDS dataset. Decitabine is a hypomethylating agent (HMA) and is approved for the treatment of patients with myelodysplastic syndromes, chronic myelomonocytic leukemia, and acute myeloid leukemia [30]. HMAs have been shown to be teratogenic, fetotoxic, and embryotoxic in preclinical studies using mice and rats [31]. There is no adequate information on the adverse effects of HMAs on human fetuses and pregnancies. However, there is a recently reported case of suspected teratogenicity of decitabine in a human fetus [30]. As with the prediction of the model, this does not completely rule out the possibility of toxicity of the drug to human fetuses. These results confirm that the model proposed in this study can predict the toxicological risk of drugs with unreported fetotoxicity in humans.

**Table 2.** False positive prediction results in the test set. Drugs labeled as ‘non-fetotoxic,’ but may not have been reported in humans. Each drug was ranked according to the prediction score given by the model. The top five drugs with prediction scores that were predicted as false positives by each model are provided.

| Rank | LR | SVM | RF | ET | GBM | XGBoost | Self-attention NN |
| --- | --- | --- | --- | --- | --- | --- | --- |
| 1 | Chloral hydrate | Diphenylpyraline | Daptomycin | Daptomycin | Decitabine | Clioquinol | Clonixin |
| 2 | Decitabine | Benzatropine | Decitabine | Decitabine | Brimonidine tartrate | Chloral hydrate | Brimonidine tartrate |
| 3 | Aminocaproic acid | Daptomycin | Brimonidine tartrate | Brimonidine tartrate | Chloral hydrate | Dexmedetomidine | Caffeine |
| 4 | Benzyl benzoate | Brimonidine tartrate | Chloral hydrate | Aminolevulinic acid | Aciclovir | Desflurane | Chloral hydrate |
| 5 | Darifenacin | Bromhexine | Benzatropine | Benzatropine | Daptomycin | Daptomycin | Desonide |

### Analysis of permutation feature importance

This study leveraged the test set to quantify the features that affect the performance of the model in terms of permutation feature importance. First, we examined the top 15 features with high importance in each machine learning model (S1 Table). Drug features that frequently appeared in the top of the feature importance list for each model are possibly highly associated with fetotoxicity. Therefore, we counted the number of appearances of each feature in the top 15 feature importance list (S2 Table). Furthermore, we examined the feature importance values estimated by each model for the important features, which were counted at least twice in the top 15 feature importance list (S1 Fig). In the result, the structural features ‘1’ and ‘80’ and the physicochemical features ALOGP and PSA had a high average feature importance. ALOGP and PSA are associated with the placental passage of the drug, which can increase the risk of potential fetotoxicity, as mentioned earlier.

Furthermore, to identify molecular substructures associated with fetotoxicity, we examined the molecular substructures represented by structural features ‘1’ and ‘80’ (Table 3). Represented by structural feature ‘1,’ quinolone derivatives, the molecular substructure of quinolone class antibiotics, are considered to have a risk of fetotoxicity [32]. Also, for mercaptobenzimidazole, fetal malformations have been reported in animal studies and attributed to maternal toxicity [33]. For benzoylhydrazine, the derivative of acetohydrazine, represented by structural feature ‘80,’ there have been reported cases of potential risk of fetotoxicity [34]. These represent the drug substructures highly associated with fetotoxicity and could be used for structural alerts. However, the number of molecular substructures represented by the ‘1’ and ‘80’ bits were 12 and 4, respectively. This result comes from the bit collision issue in Morgan fingerprints, which represent the structures as fixed-size vectors. Thus, not all substructures that can be represented by structural features highly relevant to fetotoxicity are necessarily considered as being fetotoxicity-related.

**Table 3.**
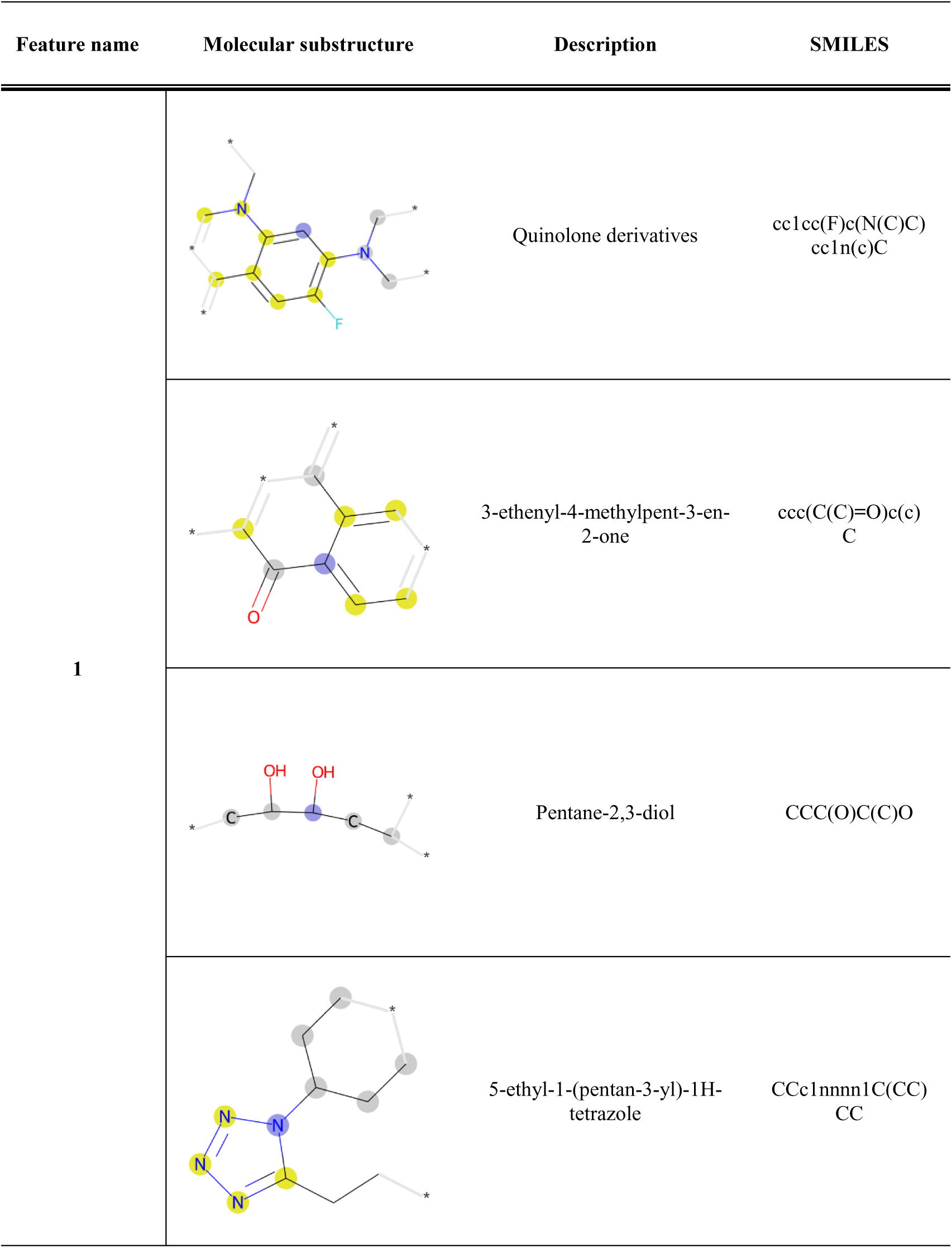
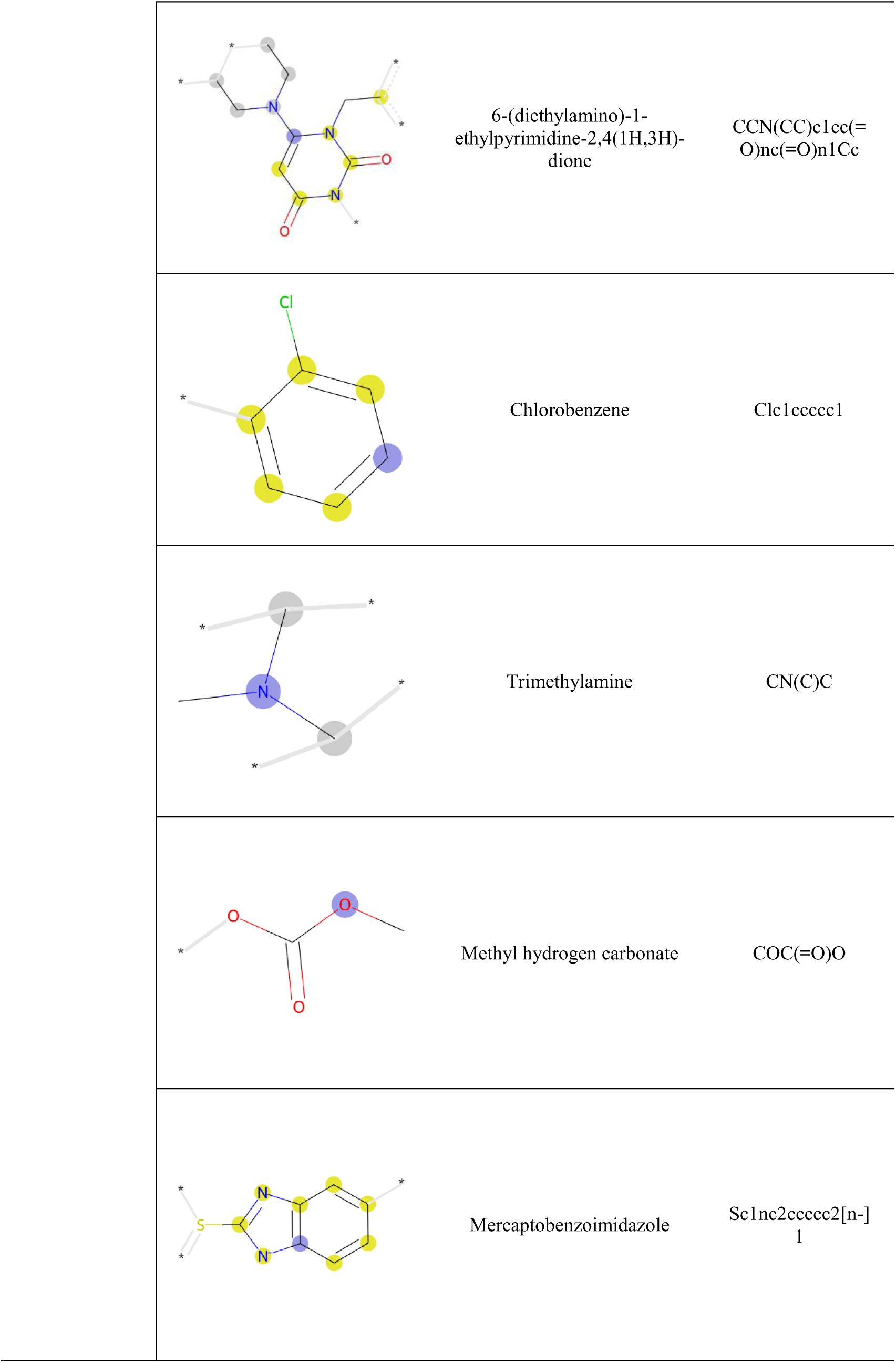
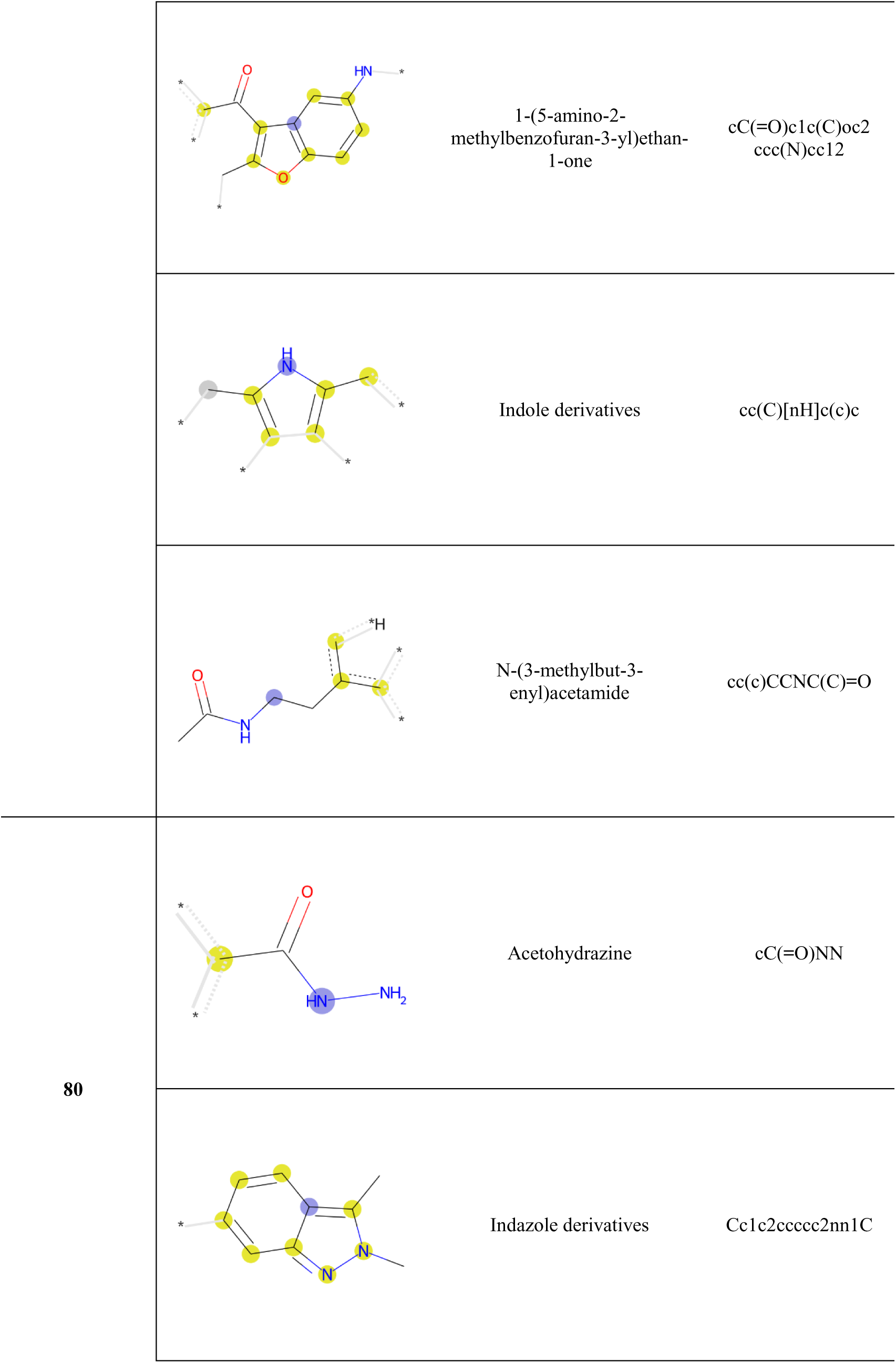
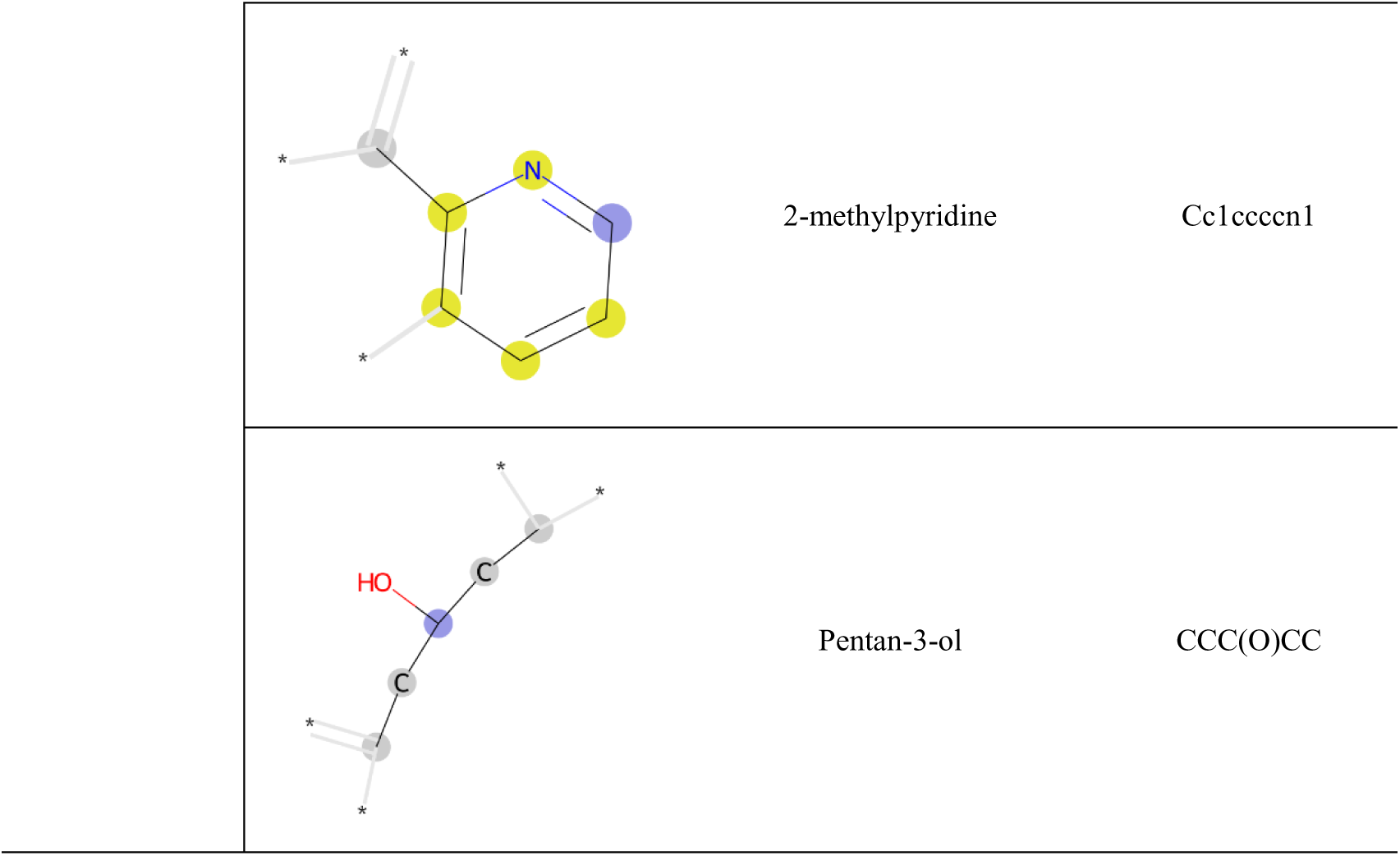
Molecular substructure corresponding to the structural feature. The molecular substructure represented by features ‘1’ and ‘80’ had a high average feature importance. For each substructure, a description and SMILES are provided.

### Analysis molecular substructure captured by attention score

This study interpreted the attention scores of each drug for which the self-attention-based NN model gave high prediction scores using the test set. For this, we highlighted and visualized the structural features of each drug with a high attention score. Then, we examined whether the highlighted structural features were related to fetotoxicity through a literature review. Fig 2 shows the visualization of the highlighted molecular substructures of the drugs with high prediction scores for fetotoxicity. Leuprorelin and nafarelin are gonadotropin-releasing hormone (GnRH) agonists (Fig 2A). GnRH is a hormone that regulates reproductive function by binding to the GnRH receptor. GnRH agonists mimic the actions of GnRH by binding to the GnRH receptor. Since continuous administration of GnRH suppresses the production of gonadotropin and ovarian steroids, long-term-acting GnRH agonists are used to treat sex steroid-dependent diseases [36]. However, a previous study has reported that GnRH agonists are potentially fetotoxic due to their mechanism of action and hormonal effects. GnRH agonists bind to the GnRH receptor, resulting in hormone regulation, and induce a hypogonadotropic-hypogonadal state, so binding affinity is key to their fetotoxicity-related mechanism of action [37]. Specifically, an Arginine residue is essential for a GnRH agonist to have a high binding affinity to the receptor [38]. Our self-attention NN model highlighted the arginine residue in leuprorelin and nafarelin. These results indicate that the model provides a molecular substructure that can explain the binding to the receptor, which is essential in the mechanism of action of GnRH agonists.

**Fig 2.**
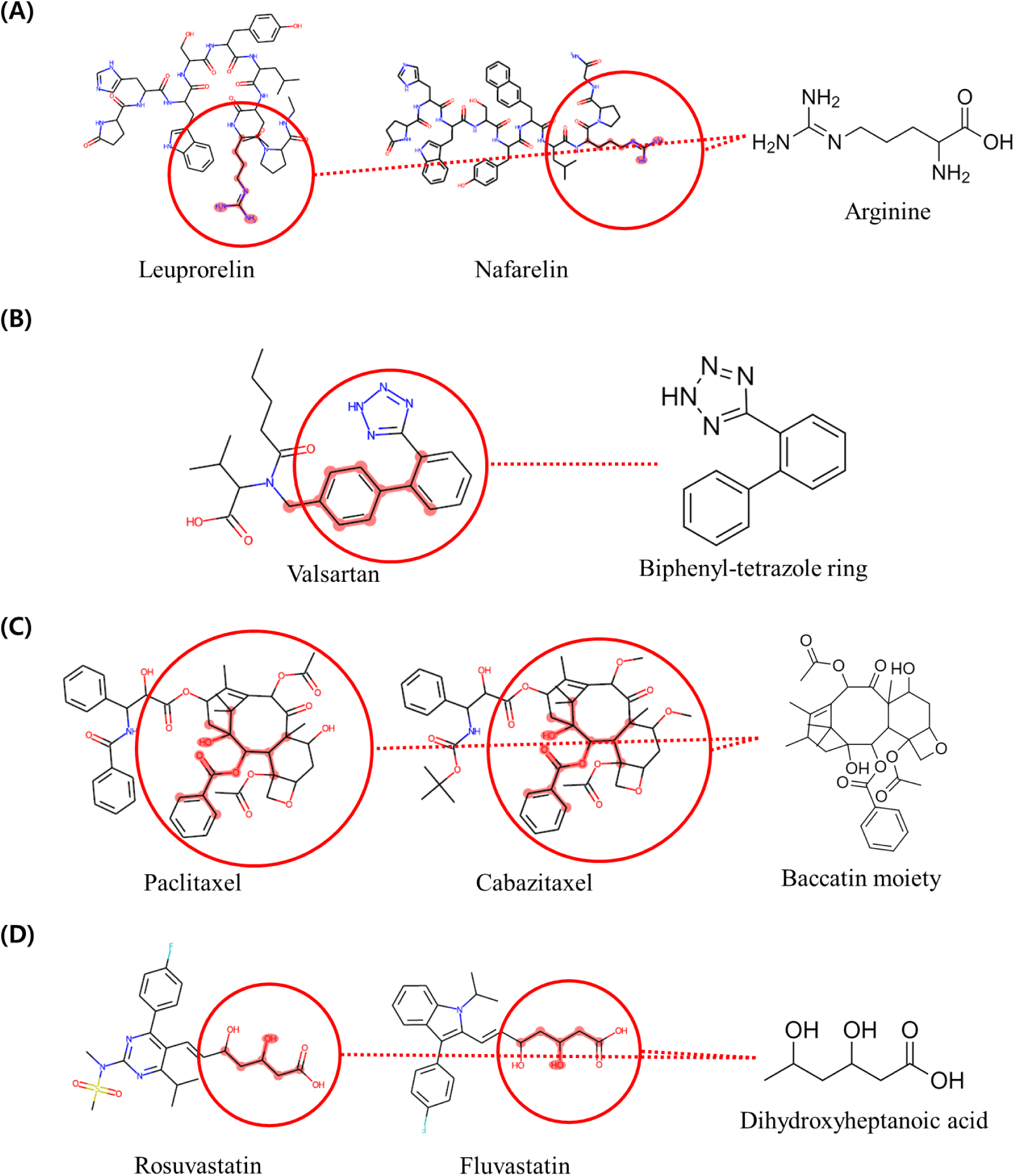
The molecular substructures of each drug that the self-attention NN model highlighted in predicting fetotoxicity. (A) The model highlighted the arginine residue of leuprorelin and nafarelin, and these drugs have high prediction scores. (B) The model gave a high attention score to the biphenyl-tetrazole ring substructure in valsartan, which has a high prediction score. (C) For the drugs paclitaxel and cabazitaxel, both with high prediction scores, the model focused on the part of baccatin moiety. (D) The model highlighted ‘dihydroxyheptanoic acid,’ the substructure of rosuvastatin and fluvastatin, which both had high prediction scores. The substructures highlighted for all seven drugs are known to be associated with fetotoxicity.

Valsartan is a drug used as an antihypertensive agent and is a type of angiotensin II receptor blocker (ARB) (Fig. 2B) [39]. ARBs reduce the action of the hormone angiotensin II, which acts directly on vascular smooth muscle cells, causing vasoconstriction and thus raising blood pressure. However, the mechanism of action of these ARBs may disrupt the fetal renin-angiotensin system, which may result in impaired fetal development [40]. The molecular substructure that most ARBs have in common is the biphenyl-tetrazole ring structure [41]. In particular, this biphenyl-tetrazole ring moiety is highly involved in binding to the angiotensin II receptor [42]. The self-attention NN model highlighted part of the biphenyl-tetrazole ring substructure of valsartan. This result confirms that our model can provide a highly relevant substructure for the mechanism of action of ARBs that may disrupt the fetal renin-angiotensin system.

The drugs cabazitaxel and paclitaxel are taxane-type anticancer drugs (Fig 2C). Taxanes are cytotoxic chemotherapeutic agents intended to inhibit tumor growth in breast, prostate, and other cancers by binding to microtubules, blocking cell cycle progression, and inducing apoptosis [43]. Cytotoxic drugs are generally contraindicated in pregnancy since they can increase the risk of teratogenic effects on the fetus [44]. The self-attention NN model focused on the ‘baccatin moiety’ of cabazitaxel and paclitaxel. This is a substructure that taxane-type anticancer drugs have in common [45]. Also, the baccatin scaffold has been suggested to be involved in holding substituents and chains in place and accurately pointing them to bind microtubules [46]. It is an important mechanism of action for taxanes that leads to cell death, the most important factor in the risk of fetotoxicity. Thus, the substructures highlighted by our model can be used to explain the mechanisms of action associated with fetotoxicity.

Rosuvastatin and fluvastatin compounds are synthetic statins that limit the synthesis of cholesterol by inhibiting hydroxymethylglutaryl coenzyme A (HMG-CoA) reductase (Fig 2D). Inhibition of cholesterol synthesis is a potential threat to fetal development, so it is contraindicated during pregnancy [47]. Synthetic statins share a common dihydroxyheptanoic acid pharmacophore moiety. Dihydroxyheptanoic acid is a moiety that mimics the HMG-CoA reductase substrate and is important in its mechanism of action in binding to HMG-CoA reductase [48]. The self-attention NN model highlighted the dihydroxyheptanoic acid, which is important in the mechanism of action associated with HMG-CoA reductase inhibition. This inhibition has been reported to be a potential threat to fetal development. Therefore, the substructure highlighted by the model can provide an explanation for the mechanism of action related to fetotoxicity. Consequently, this study confirmed that the self-attention NN model focused on significant substructures in drugs that are highly predicted to be fetotoxic.

## Discussion

In this study, the proposed fetotoxicity prediction model was interpretable with the features that are important for classification. For machine learning models, excluding the self-attention NN model, it was possible to investigate the features that are important for predicting and generating substructures. However, it was difficult to identify specific substructures associated with fetotoxicity. The permutation feature importance technique examines the features that the model depends on to make predictions, so there are limitations to its interpretation as the bits do not represent only one substructure. To overcome this, we leveraged a self-attention NN model that calculated the attention scores for each drug. Through a literature review, we confirmed that the proposed self-attention NN model could capture the specific substructures associated with fetotoxicity. Therefore, despite the highest predictive performance of the SVM and ET models, we suggest using the self-attention NN model for the benefits of transparency and interpretability.

The constructed fetotoxicity prediction model confirmed significant predictive performance and interpretability, but there are several improvements that may be considered. First, expanding the available dataset can lead to better prediction performance. The number of data points used in the study was 1,232, which might not have been sufficient to train the model. Therefore, it is necessary to consider whether reliable unstructured data, such as research literature on fetotoxicity, can be used. Second, there is the need to evaluate considerations in determining fetotoxicity, such as the route of the medications, dosage, and trimester of the pregnancy. This provides researchers with insight into the outcomes predicted by the model. Last, there is a need to consider drug feature descriptors other than the molecular fingerprint, such as the Morgan fingerprint. In this study, the structural feature vector only captured the substructure up to a radius of three for each atom. This means that the substructure of drugs larger than that was not completely captured during the interpretation of the prediction of the model. Collision problems caused by generating molecular fingerprints at a fixed vector size can also lead to obscure interpretations of model decisions. This can be particularly problematic when interpreting the common structural features of fetotoxic drugs. In this study, we set the size of the vector to 2,048 to avoid these collision problems, but this is a trade-off with the problem of trait sparsity. However, collision problems cannot be completely avoided. In the case of chlorobenzene, explained by structural feature ‘1,’ it was reported to not cause fetotoxicity at maternally toxic concentrations based on tests performed using rats and rabbits [35]. Therefore, an alternative approach to generating the vectors of structural features of drugs to overcome this challenge should be considered. These alternatives include using natural language processing models or graph neural networks to represent structural features based on the drug’s SMILES or atoms and bonds.

## Conclusion

This study proposed the FetoML model for interpretable prediction of the fetotoxicity of drugs. All constructed models had significantly high predictive performance and confirmed that they can predict the fetotoxicity of drugs. In addition, the study focused on interpretability, which is an important consideration in toxicity assessments. We used the permutation feature importance technique to identify features associated with fetotoxicity, and the results confirmed that the features were indeed associated with fetotoxicity. However, some of the substructures of the drugs represented by the features analyzed as important were not associated with fetotoxicity. Therefore, we leveraged the self-attention NN model to identify the substructures associated with fetotoxicity in each drug and examined the attention score. As a result, we confirmed that the substructures highlighted by the attention score are associated with fetotoxicity, which ensured the interpretability and transparency of the model. FetoML predicts fetotoxicity and suggests fetotoxicity-related substructures for drugs or drug candidates with less cost and time. This will improve the current paradigm of fetotoxicity testing and contribute to reducing the cost, labor, and number of animals used in testing.

## Materials and methods

### Data collection

We collected the fetotoxicity and structural information of drugs. Fetotoxicity information for the drugs was collected from the Korea Institute of Drug Safety and Risk Management (KIDS) [49], the Australian Therapeutic Goods Administration (TGA) [50], and DrugBank [51]. KIDS provides a list of drugs categorized into ‘1st class’ and ‘2nd class’ for drugs that cause or have the potential to cause fetotoxicity. This includes 1,068 drugs, and a definition of the classification and the number of drugs in each class is provided in S3 Table. The TGA provides a database that categorizes drugs into Category A, B1, B2, B3, C, D, and X according to the Australian classification system for prescribing medicines during pregnancy. This database contains fetotoxicity classification information for a total of 1,573 drugs. A detailed description of the classification criteria and the number of drugs in each category is provided in S4 Table. In addition, we collected 15 drugs from the DrugBank database that contained the term ‘teratogenic’ in the pharmacologic description of toxicity. However, in some cases, the teratogenicity information for the drugs that we collected was not reported in humans. Therefore, we manually checked and labeled them as ‘reported teratogenicity in humans’ and ‘not reported for teratogenicity in humans’ based on whether they were reported in humans. The collected drugs and their labels are provided in S5 Table. Molecular structure information of the drug was required to generate the feature vectors for input into the model. We used simplified molecular-input line-entry system (SMILES) strings to describe the molecular structure. The SMILES for each drug was collected from the Compound Combination-Oriented Natural Product Database with Unified Terminology (COCONUT) [52], PubChem [53], and ChEMBL [54].

### Data preprocessing

We preprocessed the collected data to construct a dataset that included fetotoxicity and structural information for each drug. We removed drugs with unavailable molecular structure information, such as unclear structure format and vaccines. Also, we removed cases that were not drugs, such as viruses and toxins. Subsequently, we integrated the dataset and removed drugs with duplicate SMILES strings so that only one remained. In this instance, the fetotoxicity information for the drug retained its TGA labeling. We relabeled the fetotoxicity information of the drugs in the integrated dataset according to the classification of each dataset. The classification standard description was checked for each dataset and labeled as ‘fetotoxic’ for clear risk of toxicity to human fetuses or as ‘non-fetotoxic.’ For drugs labeled with a classification in the KIDS dataset, we labeled the drugs with a ‘class 1’ classification as ‘fetotoxic’ since these have clear risk to a human fetus, and the drugs with a ‘class 2’ classification as ‘non-fetotoxic’ since these are only potentially harmful.

Similarly, the TGA dataset classifications ‘category D and X’ were labeled as ‘fetotoxic’ since these have a clear risk of harm to a human fetus, and ‘category A, B1, B2, and B3’ were labeled as ‘non-fetotoxic’ since there is insufficient evidence in human fetuses. Drugs labeled as ‘category C’ are suspected to have harmful effects on a human fetus but are not classified based on animal studies or reports in humans. Therefore, we considered there to be insufficient evidence to decide about the fetotoxicity and removed these drugs from the dataset. For drugs collected from DrugBank, we had already categorized them according to toxicity to a human fetus. Thus, we retained those labels and labeled them as ‘fetotoxic’ or ‘non-fetotoxic.’ There were 285 drugs labeled ‘fetotoxic’ and 947 drugs labeled ‘non-fetotoxic’ in the constructed dataset, for a total of 1,232 drugs. A summary of dataset construction for predicting fetotoxicity is shown in Fig 3. The constructed dataset was split into a training set for training the model and a test set for performance evaluation. To create the test set, we randomly selected 20% of the drugs classified as “fetotoxic” and randomly selected the same number of drugs classified as “non-fetotoxic.” All those drugs not included in the test set were assigned to the training set. Finally, the number of ‘fetotoxic’ drugs was 228, and the number of ‘non-fetotoxic’ drugs was 890 in the training set. The number of ‘fetotoxic’ and ‘non-fetotoxic’ drugs in the test set was 57 for each drug classification. The training set had an imbalance in the number of classes. Therefore, we reduced the class imbalance problem by applying the synthetic minority over-sampling technique (SMOTE), an oversampling technique [55].

**Fig 3.**
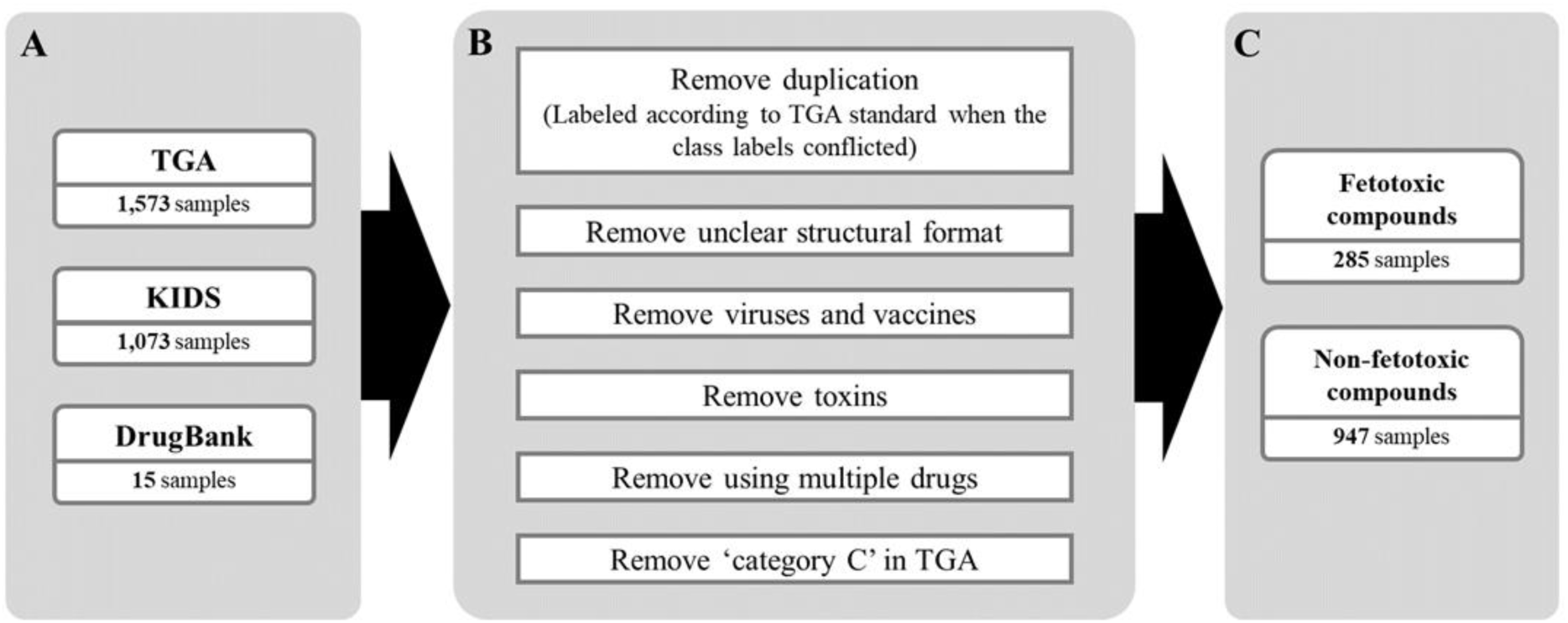
Summary of the preprocessing of the collected datasets. (A) Raw datasets on fetotoxicity were collected from TGA, KIDS, and DrugBank. (B) Preprocessing of the dataset, removing duplicate drugs, unclear structural information, and insufficient fetotoxicity information from the collected dataset. (C) The constructed dataset had a total of 1,232 drugs, with 285 labeled as ‘fetotoxic’ and 947 labeled as ‘non-fetotoxic.’

### Feature generation

In this study, feature vectors were generated for each drug to be used as the input to the model. For the representation of the drugs, structural and physicochemical features were generated based on the SMILES. We employed Morgan fingerprints to represent the environments of each atom in the molecule. Each bit in the fingerprint represented the structural features within a given radius of the atom. Therefore, it can describe the molecular substructure of a given molecule, which makes it suitable to be used as a feature vector for the interpretable model. However, molecular fingerprints are generated as a fixed-size bit vector, which can lead to bit collisions where two or more different molecular substructures are generated as the same bit. To reduce the possibility of bit collisions, we set the length of the bit vector to 2,048. Additionally, we set the radius of the atom to 3, which maximized the representation of the structural environment characteristics of each atom, enabling us to capture a more complete picture of the local molecular structure. Furthermore, we considered the physicochemical features of the drugs that are relevant to placental crossing. Certain physicochemical and molecular properties can be considered effective filters for assessing the risk of drug transfer to a fetus and potential harm to a developing fetus [56]. We focused on physicochemical features relevant to passive diffusion, the most common way drugs cross the placenta [57]. Molecular weight (MW) is a strong predictor of transfer across the placenta [58]. It is known that drugs cross the placenta when their molecular weight is less than 500 Da [59]. Based on previous studies, we assigned a value of ‘0’ to drugs with a molecular weight greater than 500 Da and a value of ‘1’ to drugs with a molecular weight less than 500 Da. In addition, it is necessary to consider the lipophilicity and polarity along with the MW of the drugs [60]. The lipophilicity and polarity of the drugs were represented by calculating AlogP and topological polar surface area (PSA), respectively. Last, we used the number of hydrogen bonding sites, which is known to be relevant for the passive diffusion of drugs [61]. The number of hydrogen bonding sites is the sum of the number of hydrogen donors and receptors present in the drug. We performed standard scaling for the AlogP, PSA, and the number of hydrogen-bonding sites, as there is no defined threshold, unlike the MW. As a result, an input vector of 2,052 features was generated for each drug, consisting of 2,048 structural features and 4 physicochemical features.

### Machine learning algorithms

LR models are generally used in various tasks that leverage input features (i.e., independent variables) from a given dataset to classify or predict dependent variables [62]. We leveraged the LR model to estimate the outcome for fetotoxicity as a probability between 0 and 1, given a vector of inputs for each drug. It can be represented by the following formula:

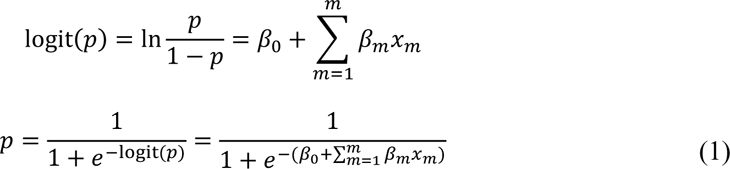

In Equation 1, *p* is the probability for fetotoxicity, logit(*p*) is the log-odds of the fetotoxicity for that drug, *β_0_* is the intercept from the regression equation, *x_m_* represents the features of the drug, *β_m_* is the regression coefficient corresponding to the features, and *m* is the number of features. Initial values of *β* were set randomly, and the value was adjusted through iterations to find the optimal coefficient that gave the best fit.

SVM is an algorithm used for classification and regression based on the statistical learning theory [63]. In this study, SVM made the optimal decision boundary according to the features of the drug for the ‘fetotoxicity’ and ‘non-fetotoxicity’ groups. However, each drug in our dataset had 2,052 dimensions of features. Since our dataset had non-linear characteristics, insufficient consideration of the kernel could have led to poor performance of the model [64]. Therefore, we used a radial basis function (RBF) kernel function, commonly used to compute the distance between each data point in a high-dimensional feature space. RBF allows the SVM algorithm to automatically determine the centers, weights, and thresholds to improve the performance of the model [65].

RF and ET are ensemble models that perform classification, regression, and other tasks by constructing multiple decision trees (DTs) trained with bootstrap aggregating (bagging). DT learns simple decision rules about data features through metrics such as Gini impurity and information gain. Although a useful model for classification, it is susceptible to bias and overfitting problems. The ensemble of multiple DTs improves these concerns and is more precise. In particular, the RF model constructed from each uncorrelated DT by combining bagging provides better performance [66]. The ET model, also known as extremely randomized trees, is a model similar to RF but with stronger randomization [67]. We constructed RF and ET models to predict fetotoxicity from the drug features.

GBM and XGBoost are ensemble models of boosting methods that perform tasks such as prediction and regression. Boosting algorithms combine weak learners (DTs) in sequential order. It weighs the data that the previous DT predicted wrong and uses the weighted data from the following DT to learn. This process is iteratively performed to generate a strong learner. GBM makes sequential weight updates as the gradient descent problem. XGBoost is a model for improving the GBM models that overfit the training dataset. It is similar to GBM but with regularization to penalize the loss as the complexity of the DT increases. In our study, we used both boosting models to construct the model to predict fetotoxicity from the features of the given drug.

The NN model is the core of deep learning algorithms and is designed to mimic biological neurons. Self-attention-based NN models leverage the self-attention mechanism with the NN model to improve performance by focusing on features that are relevant to the decision of the model for each sample during training. This allows for the interpretation of the specific features the model focuses on for making its decision. We constructed a self-attention NN model for predicting the fetotoxicity of each drug and for interpreting the highly correlated features. The structure of the model we constructed in this study is shown in Fig 4.

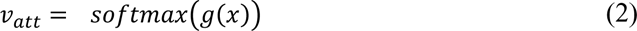

**Fig 4.**
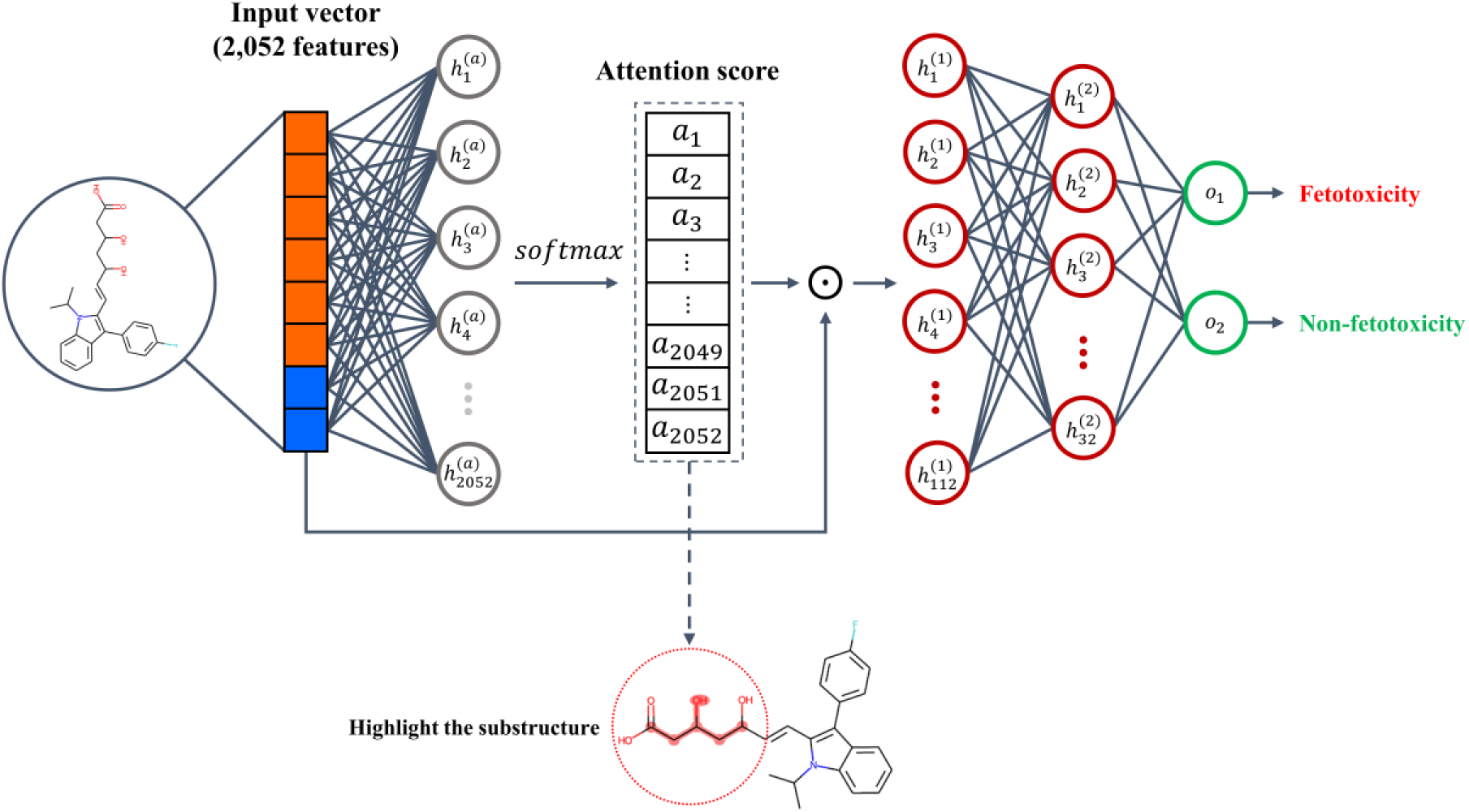
Structure of the constructed self-attention NN model. The feature vector of the drug is used as the input vector, and the number of nodes in the attention layer is equal to the number of dimensions in the input vector. Attention scores are generated through the softmax function, which performs an element-wise product with the input vector to generate weighted input values. It is then inputted into the MLP for predicting the fetotoxicity. The attention score can be interpreted as the features the model was focused on when making the predictions for each drug.

In Equation 2, *v_att_* is the attention score vector, x is the input feature vector for each drug, *g(x)* is the attention layer with no activation function, and *softmax*(*z*) is the softmax function that normalizes the elements of the vector. *g(x)* and *softmax*(*z*) can be represented by the following formula:

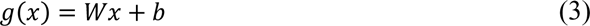

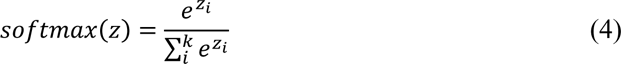

In equation 3, *x* is the input vector, *W* is the weight matrix, and *b* is the bias vector. In this study, each drug feature vector had 2,052 dimensions. The number of nodes in the attention layer was equal to the number of dimensions in the input vector. In Equation 4, *z* is the vector as an input, where *z_i_* denotes the *i_th_* element of the vector, and *k* is the number of dimensions of the vector. The softmax function normalizes the elements of the vector, so the sum of *z_i_* is 1. The obtained attention score vector performs an element-wise product with the feature vector of the drug. This can be described by the following formula:

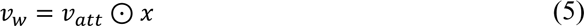

In Equation 5, ⊙ denotes the element-wise product. The learning process generates an attention score that is used to weigh the input vector, resulting in a weighted vector (*v_w_*) that highlights the relevant features for making precise predictions. The obtained weighted feature vector is inputted into the multi-layer perceptron (MLP). The hidden layers of the MLP each have the rectified linear unit (ReLU) activation function, which is computationally efficient and allows for faster convergence of the gradient descent method [68]. For each hidden layer of the MLP, batch normalization was applied [69], and the weight initialization method used was the He Normal initializer, which is effective for ReLU activation functions [70]. Additionally, we used L2 regularization to prevent the model from overfitting the training set [71, 72]. Finally, the output layer used a sigmoid activation function to generate a value between 0 and 1 as the prediction score for whether the drug is ‘fetotoxic’ or not. Since the model is a binary classification model, binary cross-entropy was used as the loss function. The optimization for training the model used the Adam optimizer.

The hyperparameters of all the constructed models were tuned using Bayesian optimization [73]. This increases the performance of the model by efficiently minimizing or maximizing the objective function for high-dimensional, nonlinear, and computationally expensive objectives. It is the exploration of hyperparameter values that can lead to the best performance for a given validation set. Using the evaluation metrics, the model was evaluated by five-fold cross-validation, and the best-performing hyperparameters were selected. The names and ranges of the optimized hyperparameters and the selected optimal values are shown in S6 Table.

The methods of the performance evaluation are described in S1 text.

### Permutation feature importance

In this study, the permutation importance algorithm was used to identify the features that were important in predicting fetotoxicity in the machine learning models. Given the trained model and the dataset as the input, the algorithm allowed for estimating the importance of features in the non-linear and opaque models [74]. The importance *i* for each feature *j* is described by the following equation:

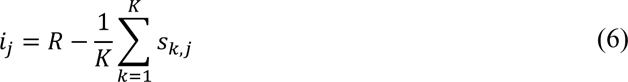

Where *R* is the reference score, which is the performance of the model when there is no random shuffling of any features, *K* is the number of repetitions. *s* is the performance of the model after randomly shuffling the features. Randomizing the values for a single feature breaks the relationship between the feature and the variables. Accordingly, the more dependent the model is on that feature, the more the performance significantly decreases, and the less dependent the model is on that feature, the less it impacts performance. This means that after randomizing the selected feature, the difference between the performance and the reference score is the quantitative indicator of the importance of the feature. In this study, the number of repetitions (*K*) was set to 30, and the evaluation metric for performance was AUROC. Furthermore, when each calculated feature importance (*i_j_*) contained zero within its 95% confidence interval, it was determined to be an invalid feature importance and was eliminated. The feature importance of each feature was then calculated by the model as the percentage of the sum of all valid feature importance. Features with high feature importance were those that were important to the model in determining the fetotoxicity of the drugs. Analyzing the features with high feature importance allowed for the identification of physicochemical features or substructures that are highly associated with fetotoxicity.

## Supporting information

### S1 text. Performance evaluation

In this study, we quantitatively evaluated the fetotoxicity prediction performance of each model. The performance was evaluated by accuracy, precision, recall, F1-score, AUROC, and AUPR. First, we used the confusion matrix to compare the predictions of the model with the actual results. From this, we calculated the number of true positives (TP), false positives (FP), true negatives (TN), and false negatives (FN). Then, we calculated the accuracy, precision, recall (TPR), F1-score, and FPR based on the obtained results. AUROC and AUPR are the areas under the curve (AUC) of the ROC and PR curves, respectively. The AUC value ranges from 0 to 1, where a higher value indicates better discrimination performance. The ROC curve plots the change in TPR and FPR as the result of the decision threshold. The y-axis of the ROC curve is TPR, and the x-axis is FPR. As the threshold increases, the TPR and FPR decrease proportionally. However, the better the classification ability of the model, the higher the TPR even at high thresholds and the lower the FPR even at low thresholds. Therefore, the more the ROC curve bends to the upper left, the better the performance, and the area under the curve increases. The PR curve for calculating the AUPR plots the change in precision and recall as the result of the decision threshold. The y-axis of the PR curve is precision, and the x-axis is recall. Precision and recall have a trade-off relationship, meaning that as one increases, the other typically decreases. Therefore, the more the PR curve bends to the upper right, the better the performance, and the area under the curve increases.

**S1 Fig.**
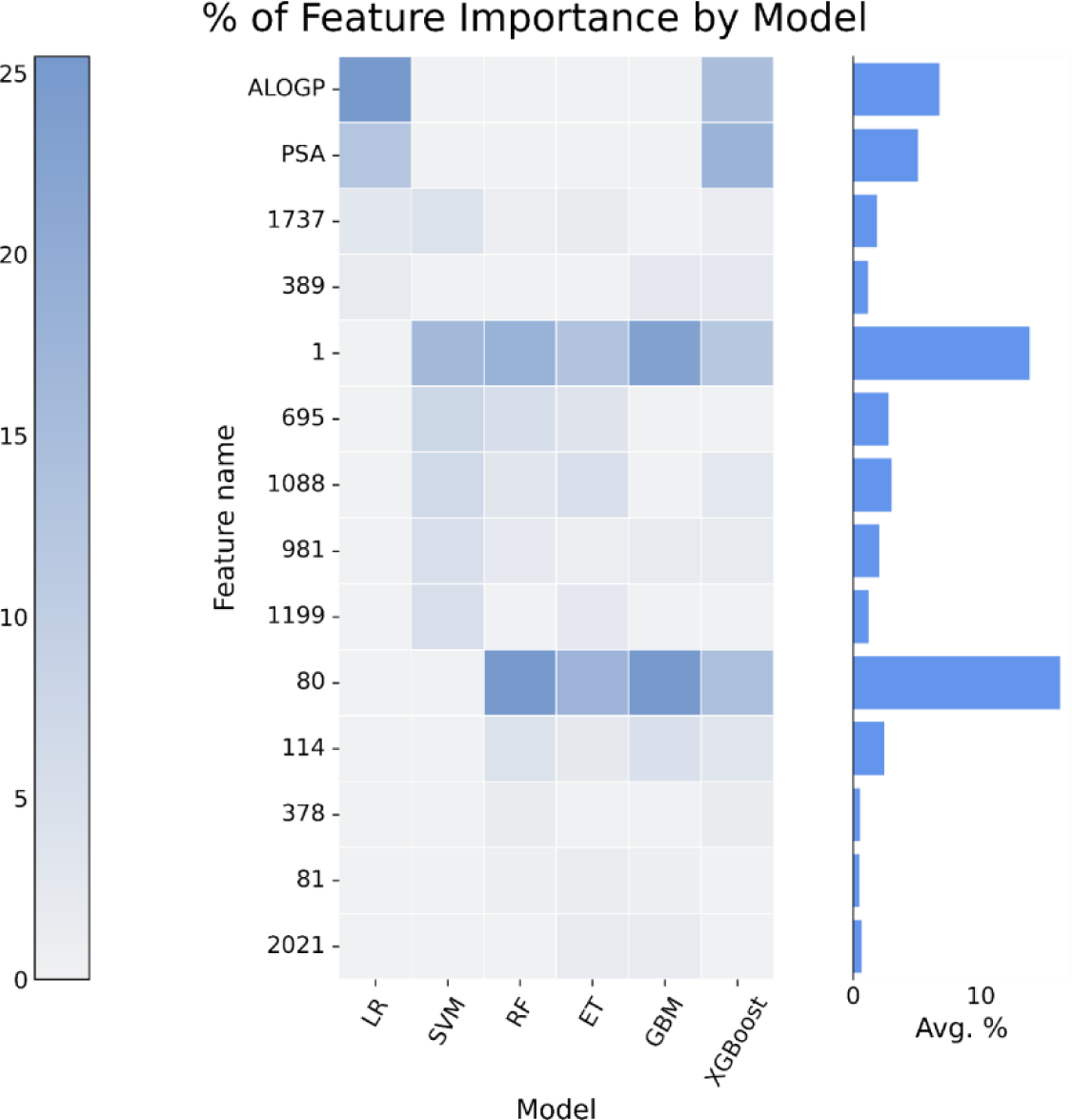
Heatmap of feature importance for each model. The x-axis is the model’s name, and the y-axis is the important feature’s name that appeared at least twice in the top feature importance list. The average over all the models for the given feature is on the right.

**S1 Table.** Permutation feature importance results for the machine learning models, including the LR, SVM, RF, ET, GBM, and XGBoost models. Each feature was ranked by the importance of the feature in each machine learning model, and the top 15 features are shown.

| Rank | LR | SVM | RF | ET | GBM | XGBoost |
| --- | --- | --- | --- | --- | --- | --- |
| 1 | ALOGP | 1 | 80 | 80 | 80 | PSA |
| 2 | PSA | 695 | 1 | 1 | 1 | ALOGP |
| 3 | 1737 | 1088 | 1873 | 1873 | 1550 | 80 |
| 4 | 941 | 981 | 695 | 695 | 114 | 1 |
| 5 | 693 | 119 | 114 | 114 | 1573 | 807 |
| 6 | 67 | 1199 | 1088 | 1088 | 389 | 114 |
| 7 | 537 | 1737 | 227 | 227 | 41 | 1088 |
| **8** | 1701 | 1932 | 981 | 981 | 1760 | 389 |
| **9** | 389 | 387 | 378 | 378 | 2021 | 981 |
| **10** | 1908 | 803 | 1849 | 1849 | 981 | 1449 |
| **11** | 1499 | 175 | 1737 | 1737 | 2043 | 1866 |
| **12** | 1626 | 902 | 456 | 456 | 81 | 378 |
| **13** | 1880 | 1839 | 1581 | 1581 | 607 | 926 |
| **14** | 1411 | 1757 | 1351 | 1351 | 1698 | 1737 |
| **15** | 995 | 1014 | 81 | 81 | 1860 | 564 |

**S2 Table.** The count of features that appeared at least twice in the top 15 feature importance list. Each feature was ranked according to its count.

| **Rank** | **Feature name** | **Count** |
| --- | --- | --- |
| **1** | 1 | 5 |
| **2** | 1737 | 5 |
| **3** | 1088 | 4 |
| **4** | 114 | 4 |
| **5** | 80 | 4 |
| **6** | 981 | 4 |
| **7** | 389 | 3 |
| **8** | 695 | 3 |
| **9** | 378 | 2 |
| **10** | 81 | 2 |
| **11** | PSA | 2 |
| **12** | 2021 | 2 |
| **13** | 1199 | 2 |
| 14 | ALGOP | 2 |

**S3 Table.**
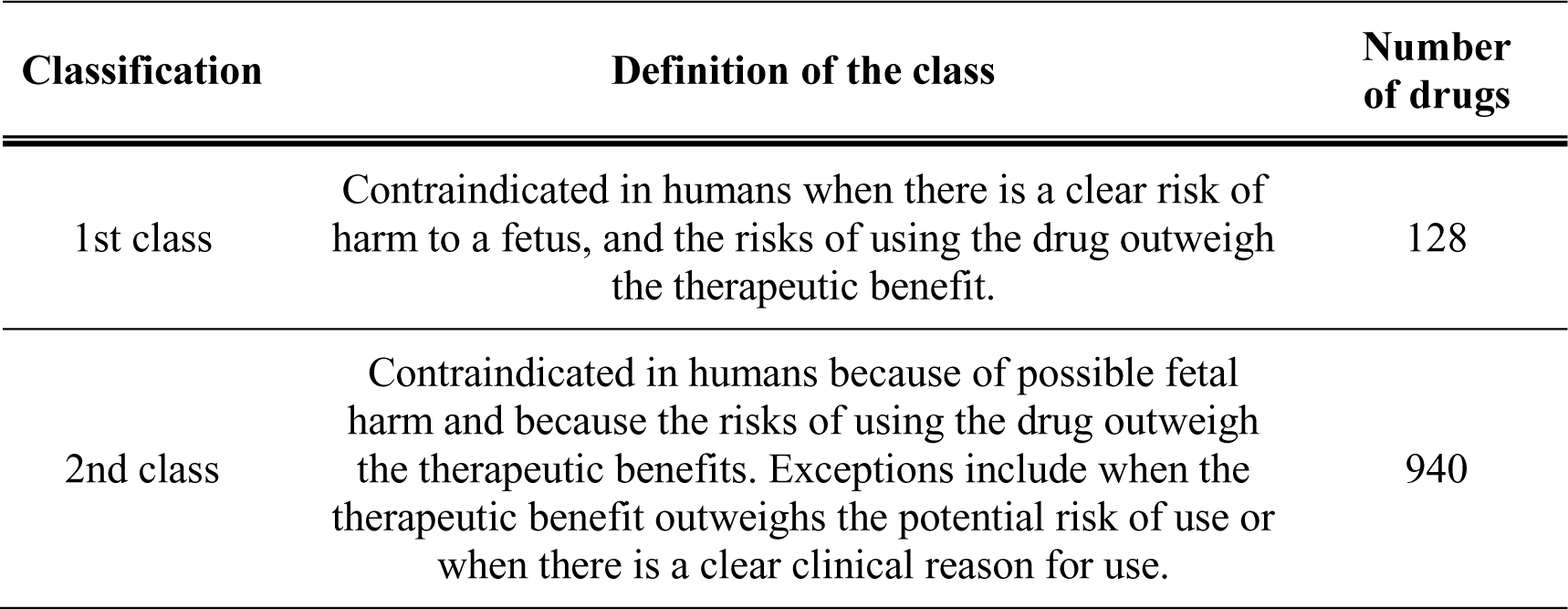
The composition of the KIDS dataset and descriptions of each classification label. Drugs classified as ‘1st class’ pose a clear risk of toxicity to a human fetus, while drugs classified as ‘2nd class’ pose only the possibility of risks to a human fetus.

**S4 Table.**
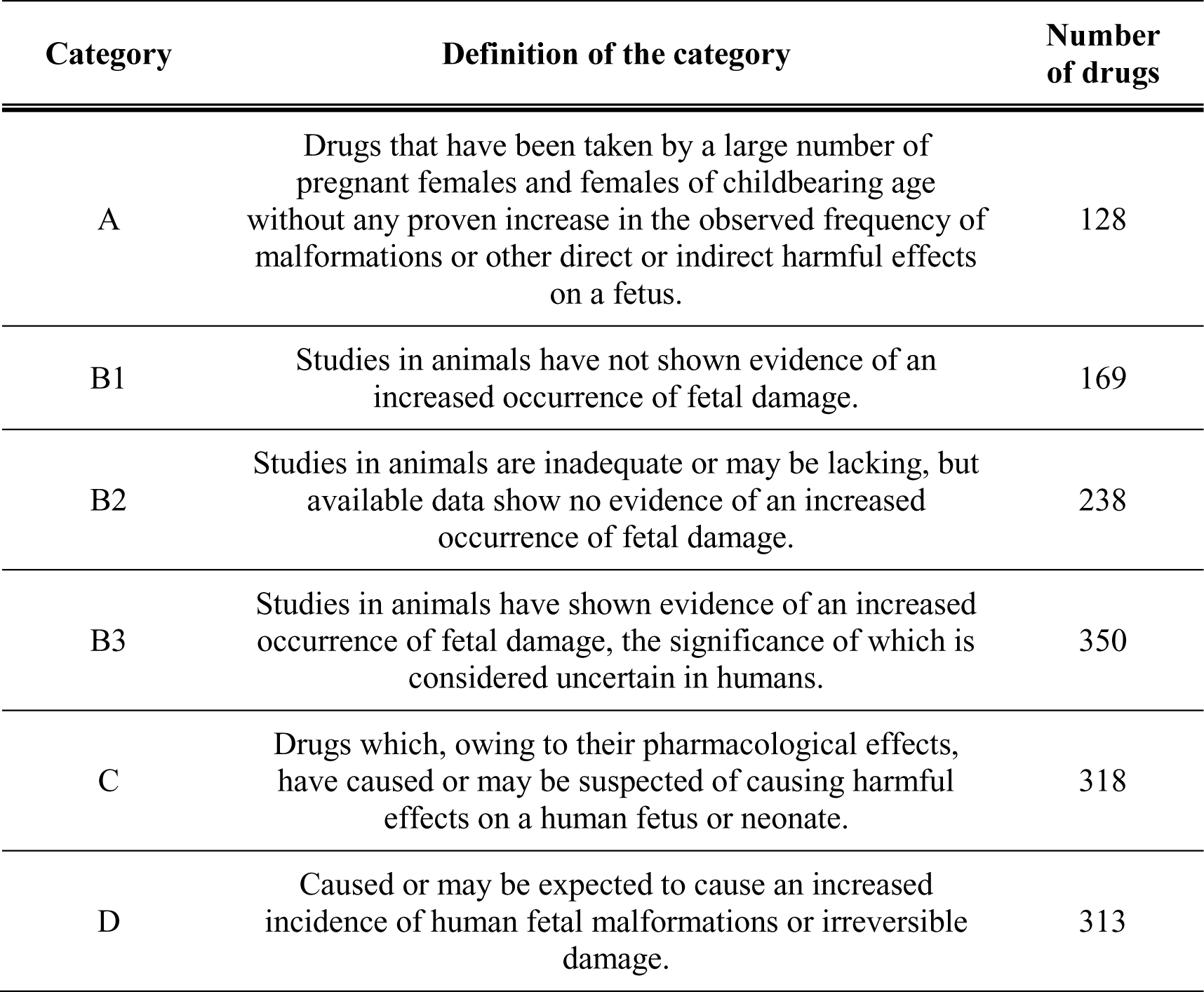
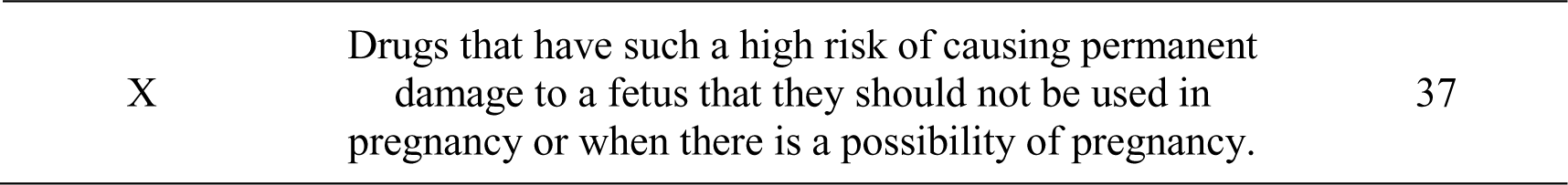
The composition of the TGA dataset and descriptions of each classification label. ‘Category A, B1, B2, and B3’ drugs have no reported fetotoxicity in humans; ‘Category C’ drugs have insufficient experimental results but pose a risk of fetotoxicity from their pharmacologic effects; and ‘Category D and X’ drugs have reported fetotoxicity in humans.

**S5 Table.** List of drugs collected from the DrugBank dataset and each label. Each drug was manually reviewed for toxicity descriptions and categorized according to whether it posed a clear fetotoxicity risk in humans.

| Drug name | Classification |
| --- | --- |
| fluticasone propionate | Not reported for teratogenicity in humans |
| tazarotene | Not reported for teratogenicity in humans |
| d-amphetamine | Teratogenicity reported in humans |
| docosahexaenoic acid | Teratogenicity reported in humans |
| ambrisentan | Teratogenicity reported in humans |
| fluticasone furoate | Not reported for teratogenicity in humans |
| olaparib | Not reported for teratogenicity in humans |
| lenvatinib | Teratogenicity reported in humans |
| norprolac | Not reported for teratogenicity in humans |
| eslicarbazepine acetate | Not reported for teratogenicity in humans |
| benidipine hydrochloride | Not reported for teratogenicity in humans |
| dextran | Not reported for teratogenicity in humans |
| 10-undecenoic acid | Not reported for teratogenicity in humans |
| DHEC | Not reported for teratogenicity in humans |
| fluticasone | Not reported for teratogenicity in humans |

**S6 Table.** List of hyperparameters tuned by using Bayesian optimization. The name of each model’s tuned hyperparameter, the ranges that were examined, and the optimal values that were estimated to have the best performance are provided.

| Model | Hyperparameter name | Range | Optimal value |
| --- | --- | --- | --- |
| LR | C | [0.001, 100] | 0.27 |
| SVM | C | [0.001, 100] | 7.92 |
|  | gamma | [0.0001, 0.1] | 0.035 |
| RF | n_estimators | [10, 250] | 90 |
|  | min_samples_split | [2, 25] | 5 |
|  | max_features | [0.1, 0.999] | 0.34 |
|  | max_depth | [30, 100] | 203 |
|  | min_samples_leaf | [1, 10] | 1 |
| ET | n_estimators | [10, 250] | 34 |
|  | min_samples_split | [10, 50] | 43 |
|  | max_features | [0.1, 0.999] | 0.10 |
|  | max_depth | [20, 100] | 24.09 |
|  | min_samples_leaf | [1, 20] | 1 |
| GBM | learning rate | [0.001, 0.1] | 0.063 |
|  | n_estimators | [200, 500] | 246 |
|  | max_depth | [1, 100] | 93 |
|  | min_samples_split | [2, 25] | 5 |
|  | min_samples_leaf | [1, 25] | 2 |
|  | subsample | [0, 1] | 0.85 |
| XGBoost | eta | [0.001, 0.1] | 0.035 |
|  | n_estimators | [300, 700] | 459 |
|  | min_child_weight | [1, 2] | 1 |
|  | gamma | [0, 2] | 1.18 |
|  | max_depth | [5, 15] | 6 |
|  | subsample | [0.5, 1] | 0.87 |
|  | colsample_bytree | [0.5, 1] | 0.94 |
| Self-attention NN | learning rate | [0.001, 0.01] | 0.003 |
|  | Num. of layers | [1, 2, 3] | 2 |
|  | Num. of nodes (1st layer) | [4, 128] | 116 |
|  | Num. of nodes (2nd, 3rd layer) | [4, 32] | 32 |

## Notes

### Competing Interest Statement

The authors have declared no competing interest.

